# Childhood Trauma and Transdiagnostic Mental Health Outcomes Among Adults in 17 Arab Countries: A Multinational Cross-Sectional Study Using a Latent Global Trauma Model

**DOI:** 10.64898/2026.09.19.752885

**Authors:** Khalid Mohammed Al-Dhayani, Baraa Abdelazim Osman, Doaa Magzub Elrayah, Yasmin Elsayed Hassan, Ayah Nael Abusafyeh, Serene El Fil, Miaad Said Almazrouai, Baneen Hashim Hani, Yasmine Noamen Rebai, Amna Said Almazroei, AlRafa Salah Alkhanbashi, Mahdi W. Suboh, Nareman Ali Qari, Abdulaziz Abdullah Alsulqi, Lahreche Silouane Khadidja

## Abstract

**Background:** Childhood trauma is a well-established risk factor for adverse mental health outcomes, yet large-scale cross-national evidence from the Arab region remains limited. This study examined the prevalence of childhood trauma and its associations with multiple transdiagnostic outcomes across 17 Arab countries.

**Methods:** A cross-sectional online survey was conducted between September and October 2025 among 6,181 adults from 17 Arab countries. Childhood trauma was assessed using the Childhood Trauma Questionnaire–Short Form (CTQ-SF). Transdiagnostic mental health outcomes included depression, anxiety, posttraumatic stress symptoms (PTSS), maladaptive behavior, and social/functional impairment. Descriptive statistics, Pearson correlations, and multiple linear regression analyses were performed via IBM SPSS Statistics. Confirmatory factor analysis and structural equation modeling were conducted via R (Lavaan) to evaluate a second-order latent construct representing global childhood trauma.

**Results:** Childhood trauma was widely reported across the sample, with physical neglect and emotional neglect emerging as the most frequently endorsed domains. All trauma domains were significantly associated with poorer mental health outcomes (p < .001). In regression analyses adjusted for demographic factors, emotional abuse emerged as the strongest predictor of depression (β = .472, p < .001) and PTSS (β =.439, p < .001). Structural equation modeling supported a second-order global trauma construct, which showed strong associations with depression (β = .70), anxiety (β = .64), PTSS (β = .68), maladaptive behavior (β = .68), and social impairment (β = .67). This latent factor showed strong associations with all mental health outcomes, explaining substantial variance in depression (R² = .52), anxiety (R² = .43), and PTSS (R² = .48). The structural model demonstrated borderline-to-acceptable fit (CFI = .899, TLI = .890, RMSEA = .058, SRMR = .085), supporting the hypothesized latent structure while indicating scope for further model refinement.

**Conclusions:** Childhood trauma is highly prevalent across the Arab region and is strongly associated with multiple dimensions of psychological distress, highlighting the urgent need for trauma-informed prevention, screening, and early intervention programs in the Arab region.

## 1. Introduction

Childhood is a critical period for lifelong psychological and physiological development. Exposure to trauma and adversity—often conceptualized as adverse childhood experiences (ACEs)—has been consistently linked to altered neurodevelopment and long-term health burdens.^1^ According to the World Health Organization, up to one billion children experience violence or neglect annually, and one in seven adolescents lives with a mental disorder.^2, 12^ These early adversities extend beyond individual suffering, contributing to substantial societal costs through increased healthcare utilization, reduced productivity, and intergenerational transmission of risk.^3–5^

Childhood trauma is associated with long-term alterations in emotional and cognitive functioning, manifesting alongside a wide range of psychiatric disorders. These include depression, anxiety disorders, posttraumatic stress disorder (PTSD), and substance use disorders.^6-9^ Exposure to violence or neglect during childhood is linked to overwhelmed emotional regulatory capacities and increased vulnerability to psychopathology later in life.^10^ Individuals exposed to early trauma may adopt maladaptive coping strategies, including substance use, to manage emotional dysregulation.^9^

Empirical evidence from large-scale studies indicates that cumulative exposure to childhood trauma is significantly associated with a greater likelihood of adverse adult outcomes,^11^ including PTSD, alcohol dependence, injection drug use, tobacco use, involvement in sex work, chronic medical conditions, and poor overall quality of life.^11,12^ Collectively, these findings consistently demonstrate a robust association between childhood trauma and adverse adult transdiagnostic psychopathology.

The Middle East and North Africa (MENA) region presents a uniquely complex context for examining the long-term correlates of childhood trauma. Children in this region are disproportionately exposed to psychological and physical harm due to armed conflict, political instability, socioeconomic deprivation, and displacement relative to global averages.^13–15^ Despite this elevated risk, pervasive stigma surrounding mental illness frequently suppresses disclosure of traumatic experiences and limits access to mental health services.^16,17^

Empirical evidence indicates high rates of childhood maltreatment across MENA populations, including physical abuse, emotional neglect, and exposure to domestic and community violence.^18–20^ In Oman, 88% of adults reported at least one ACE, with 38.2% reporting four or more.^21^ In Abu Dhabi, the United Arab Emirates, respondents reported an average of 1.74 ACEs.^22^ Yemen presents similarly alarming rates, with child abuse affecting an estimated 51–81% of children across homes, schools, and juvenile institutions.^23^ These contextual factors highlight the importance of examining childhood trauma within the MENA region. However, most existing research has been conducted in Western populations, limiting generalizability to Arab societies. Distinct sociocultural norms, political instability, armed conflict, displacement, and stigma surrounding mental illness are related to a region’s mental health landscape in unique ways. Nevertheless, multicountry research on childhood trauma in Arab societies remains scarce. Existing regional studies have reported prevalence rates, including 40% in Sudan,^24^ 22.1% in Qatar,^25^ and 47.26% in Bahrain.^26^ The present study aims to address this critical research gap by evaluating the prevalence and psychological correlates of childhood trauma across 17 Arab nations.

A second-order latent modeling approach was selected because childhood trauma domains are conceptually and empirically interrelated, allowing estimation of the shared underlying burden of adversity while accounting for measurement error.^27,31^ This framework provides a theoretically meaningful representation of cumulative trauma that is more informative than simple composite scores and better aligned with latent variable theory than approaches such as network analysis, which focus on relationships among observed variables rather than a common underlying construct.^43,44^ Previous psychometric investigations have also shown that reverse-coded neglect items within the Childhood Trauma Questionnaire–Short Form (CTQ-SF) may exhibit atypical measurement behavior, particularly in multilingual and cross-cultural settings, underscoring the importance of carefully evaluating their performance in multinational samples.^28,45^

Accordingly, the present study tested two a priori hypotheses. First, we hypothesized that the five CTQ-SF domains (emotional abuse, physical abuse, sexual abuse, emotional neglect, and physical neglect) would load significantly onto a higher-order latent “Global Trauma” construct. Second, we hypothesized that this latent Global Trauma construct would significantly predict greater severity of depression, anxiety, posttraumatic stress symptoms (PTSS), maladaptive behavior, and social/functional impairment among adults across the 17 participating Arab countries.

## 2. Methods

### 2.1 Study Design

This study employed a multinational, descriptive, cross-sectional survey design to examine the associations between adverse childhood experiences and adult transdiagnostic psychiatric symptom domains across 17 Arab countries. The manuscript was prepared in accordance with the STROBE 2021 reporting guidelines for cross-sectional studies. The study was conducted in accordance with the ethical principles of the Declaration of Helsinki. As Amran University does not currently have an established Institutional Review Board, formal IRB approval was not available. Participation was voluntary and anonymous, electronic informed consent was obtained from all participants, and no personally identifiable information was collected.

### 2.2 Setting and Participants

Data were collected from a large sample of adults residing in 17 Arab countries: Jordan, the United Arab Emirates, Bahrain, Algeria, Saudi Arabia, Sudan, Iraq, Morocco, Yemen, Tunisia, Syria, Oman, Palestine, Qatar, Lebanon, Libya, and Egypt. Data collection was conducted between September 1, 2025, and October 30, 2025. Although country representation was relatively balanced, the sample was heavily skewed toward young, highly educated female participants recruited via social media, which reflects typical participation patterns in large-scale online surveys. Recruitment was conducted through online platforms, including Facebook, Instagram, WhatsApp, and university-based social networks, using voluntary participation invitations shared by the research team and collaborating networks. No paid advertisements or targeted recruitment campaigns were used. As a convenience online sample, this approach may have introduced selection bias by preferentially reaching younger, urban, educated, and digitally connected individuals, while potentially underrepresenting older adults, rural populations, individuals with lower digital literacy, and those without reliable internet access. Therefore, the findings should be interpreted as reflecting the experiences of participating adults rather than as nationally representative estimates for the general populations of the 17 Arab countries. The eligibility criteria included being between 18 and 55 years of age, having a current residence in one of the participating countries, and providing electronic informed consent. Responses were excluded if trauma or symptom data were incomplete (>20% missing responses) or if duplicate submissions were identified. To ensure data integrity, IP address filtering and “limit to one response” settings were applied within the survey platform.

### 2.3 Procedures and Measures

The survey was administered via Google Forms. The CTQ-SF and mental health outcome items were translated using a forward–backward translation procedure. A single standardized Arabic version was used across the participating countries to ensure consistency, with linguistic review performed by bilingual researchers familiar with Arabic-speaking populations. Because Arabic-speaking countries share a common written language while differing in dialects, minor wording adjustments were reviewed during pilot testing to ensure cultural comprehensibility without altering the meaning of the original items. All instruments underwent forward– backward translation and pilot testing with 50 participants to ensure linguistic and cultural validity.

The questionnaire consisted of three components (Supplementary Appendix A): (1) sociodemographic characteristics, including gender, age, education level, occupation, and country of residence; (2) childhood trauma exposure measured using the 28-item Childhood Trauma Questionnaire–Short Form (CTQ-SF),^27,28^ which assesses emotional abuse, physical abuse, sexual abuse, emotional neglect, and physical neglect; and (3) current psychological symptoms assessed using a 22-item Mental Health Outcome Questionnaire capturing transdiagnostic domains including depressive symptoms, anxiety symptoms, posttraumatic stress symptoms (PTSS), maladaptive behavior, and social/functional impairment. The questionnaire was developed on the basis of DSM-5 symptom domains and previously used transdiagnostic symptom frameworks.^29,30^ All items were rated on a 5-point Likert scale reflecting frequency or agreement with each statement. Crucially, positively worded neglect items (e.g., assessing the presence of love, support, or adequate care) were reverse-coded prior to analysis. Because of this reverse coding, higher subscale scores for both emotional neglect and physical neglect consistently reflect a greater severity of neglect.

### 2.4 Statistical analysis

All the statistical analyses were conducted via IBM SPSS Statistics (version 29.0) and R (lavaan package). Descriptive statistics were calculated to summarize the demographic characteristics and prevalence of childhood trauma domains. The internal consistency of all scales was evaluated via Cronbach’s α coefficients. Distributional assumptions were examined; given the large sample size (N = 6,181), parametric analyses were applied under the Central Limit Theorem.

Pearson product–moment correlation coefficients were computed to examine bivariate associations between childhood trauma domains and mental health outcomes. Separate multiple linear regression models were estimated to evaluate the predictive effects of the five trauma domains (emotional abuse, physical abuse, sexual abuse, emotional neglect, and physical neglect) on depression, anxiety, Posttraumatic Stress Symptoms (PTSS), maladaptive behavior, and social impairment, adjusting for demographic covariates (age, sex, and education). Regression assumptions, including linearity, homoscedasticity, independence of errors, and multicollinearity, were examined and met; variance inflation factors ranged from 1.18-2.34, indicating no concerns of multicollinearity.

To examine the latent structure of childhood trauma and its global association with mental health outcomes, confirmatory factor analysis (CFA) and structural equation modeling (SEM) were conducted via the *lavaan* package in R. A second-order latent construct comprising the five trauma domains was specified based on the theoretical assumption that different forms of childhood adversity represent interconnected manifestations of an overarching global trauma burden. This approach allows estimation of the shared variance among trauma domains while retaining the contribution of each specific trauma domain. Although alternative models, such as bifactor models, may provide additional information by separating general trauma variance from domain-specific variance and potential wording-related method effects, the present study focused on testing the higher-order latent trauma framework. Model fit was evaluated using the CFI, TLI, RMSEA, and SRMR according to established guidelines. The final model demonstrated borderline acceptable fit (CFI = 0.899, TLI = 0.890, RMSEA = 0.058, SRMR = 0.085), and these indices were interpreted cautiously rather than as evidence of optimal model fit. Four items (soc4, sa15, pn23, and pn24) were removed after examination of factor loadings and modification indices indicated poor psychometric performance. These modifications were conducted to improve measurement adequacy while preserving the theoretical structure of the CTQ-SF domains. Because each trauma domain remained represented after item removal, the conceptual meaning of the latent constructs was considered preserved; however, the possibility of cultural interpretation and reverse-wording effects influencing item performance was acknowledged.

Multigroup Confirmatory Factor Analysis (MGCFA) was additionally conducted to examine measurement invariance across gender and country groups. Statistical significance was set at p < .05 (two-tailed).

## 3. Results

### 3.1 Sociodemographic Characteristics

The final analytic sample consisted of 6,181 adults from 17 Arab countries. Sociodemographic characteristics of the participants are presented in Table 1. The sample was predominantly female (68.8%), while 31.2% were male. Most respondents were young adults aged 18–25 years (70.6%), followed by those aged 26–35 years (18.7%). Most participants held a bachelor’s degree (55.2%). Country representation was relatively balanced across the 17 participating nations.

**Table 1.** Sociodemographic Characteristics of the Study Participants (N = 6,181)

| Variable | Category | n | % |
| --- | --- | --- | --- |
| **Age group (years)** | 18–25 | 4,366 | 70.6 |
|  | 26–35 | 1,155 | 18.7 |
|  | 36–45 | 434 | 7.0 |
|  | 46–55 | 226 | 3.7 |
| **Gender** | Male | 1,931 | 31.2 |
|  | Female | 4,250 | 68.8 |
| **Education level** | Uneducated | 67 | 1.1 |
|  | High school | 1,095 | 17.7 |
|  | Diploma | 660 | 10.7 |
|  | Bachelor | 3,412 | 55.2 |
|  | Postgraduate | 946 | 15.3 |
| **Occupation** | Unemployed | 513 | 8.3 |
|  | Student | 3,706 | 60.0 |
|  | Freelancer/Self-employed | 323 | 5.2 |
|  | Retired | 77 | 1.2 |
|  | Private sector | 529 | 8.6 |
|  | Public sector | 355 | 5.7 |
|  | Health practitioner | 678 | 11.0 |
| **Country of residency** | Jordan | 372 | 6.0 |
|  | UAE | 368 | 6.0 |
|  | Bahrain | 380 | 6.1 |
|  | Algeria | 370 | 6.0 |
|  | Saudi Arabia | 356 | 5.8 |
|  | Sudan | 351 | 5.7 |
|  | Iraq | 381 | 6.2 |
|  | Morocco | 370 | 6.0 |
|  | Yemen | 373 | 6.0 |
|  | Tunisia | 370 | 6.0 |
|  | Syria | 342 | 5.5 |
|  | Oman | 373 | 6.0 |
|  | Palestine | 347 | 5.6 |
|  | Qatar | 368 | 6.0 |
|  | Lebanon | 358 | 5.8 |
|  | Libya | 350 | 5.7 |
|  | Egypt | 352 | 5.7 |

**Note:** *n* = number of participants; % = percentage of the total sample.

### 3.2 Psychometric Validation and Measurement Model

Prior to structural modeling, the psychometric properties of the study instruments were evaluated using confirmatory factor analysis (CFA). As shown in Table 2, the measurement model demonstrated **borderline** acceptable overall fit (χ²(954) = 20,574.74, CFI = 0.899, TLI = 0.890, RMSEA = 0.058, SRMR = 0.085).

**Table 2.**
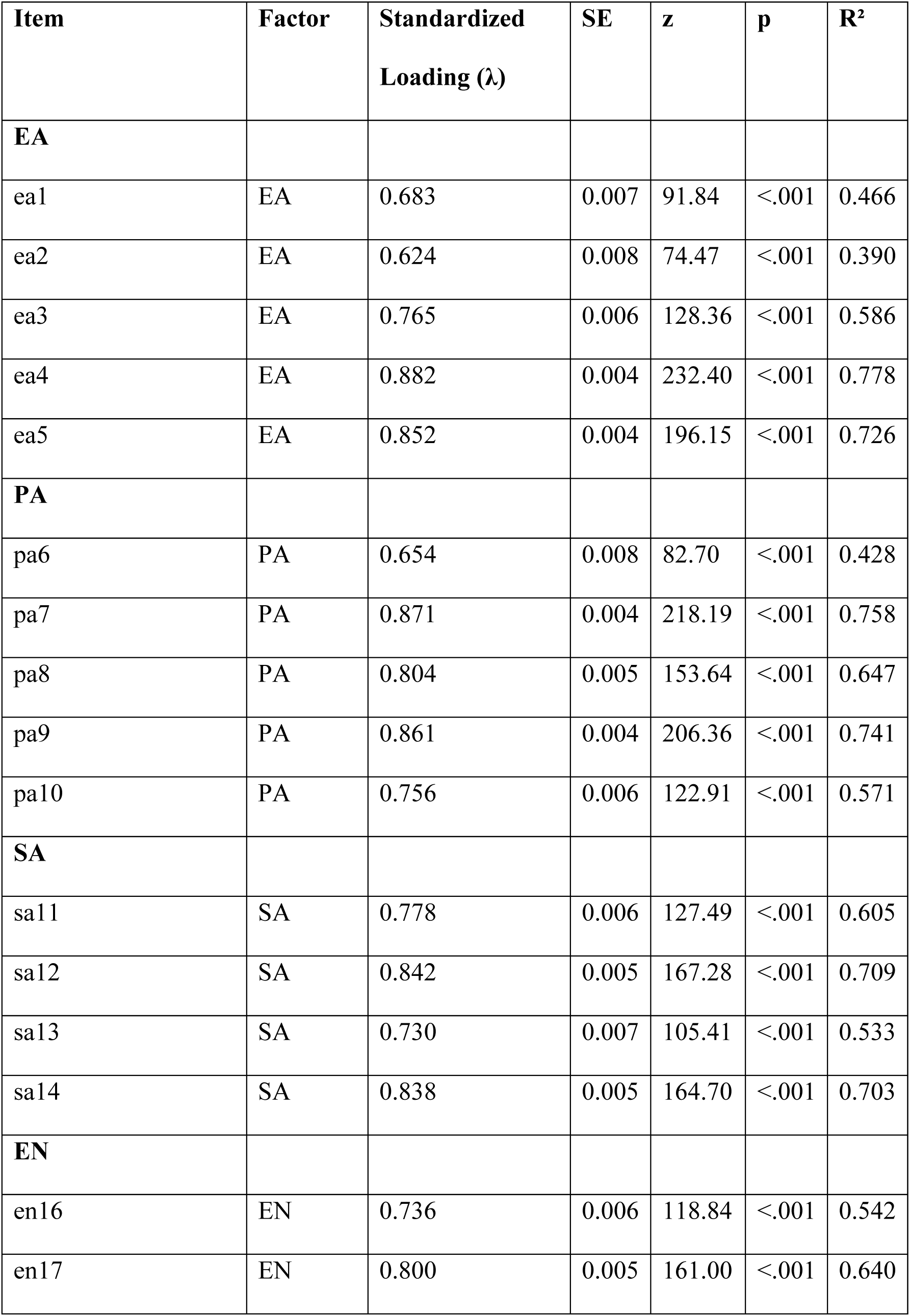
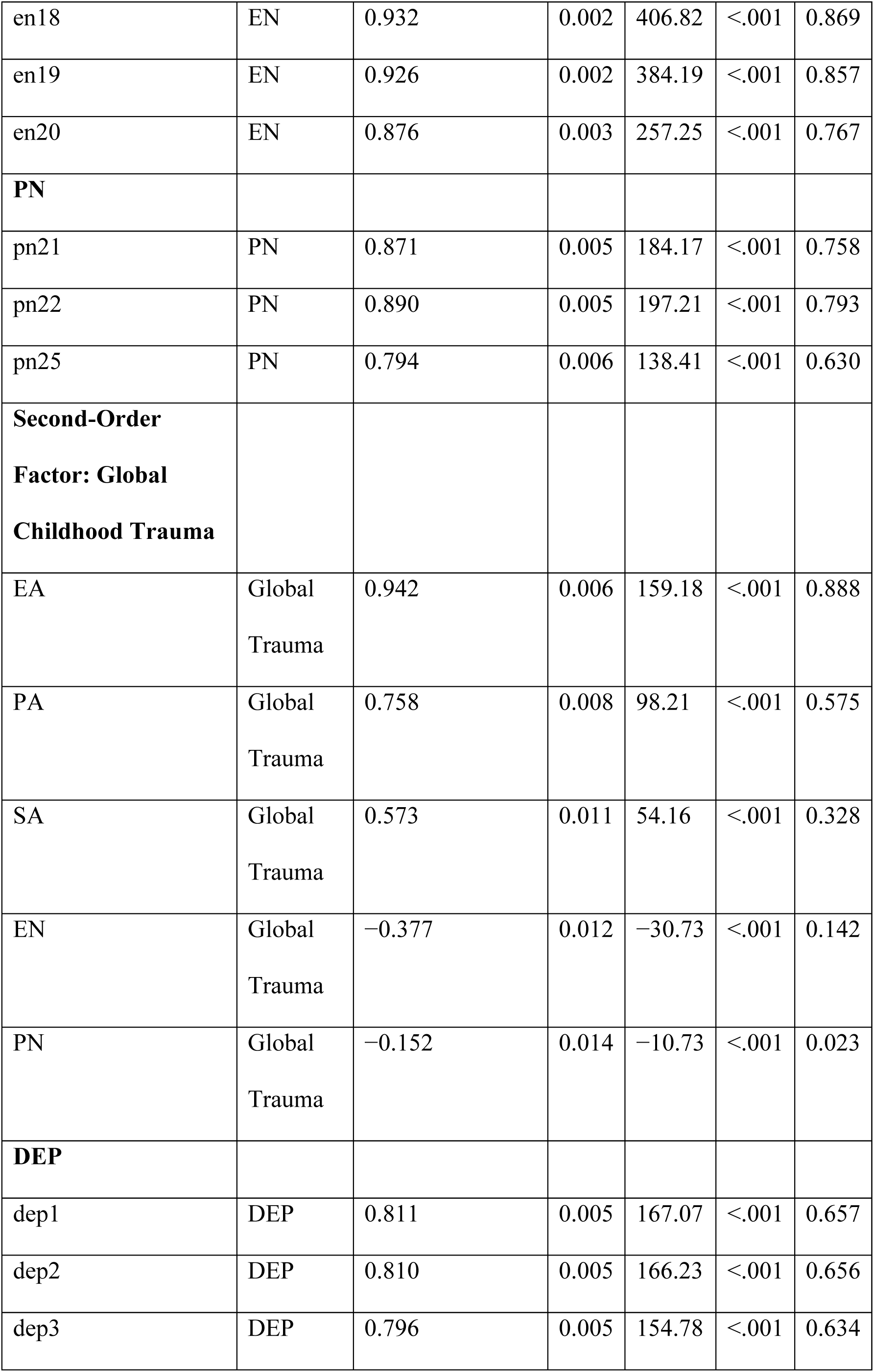
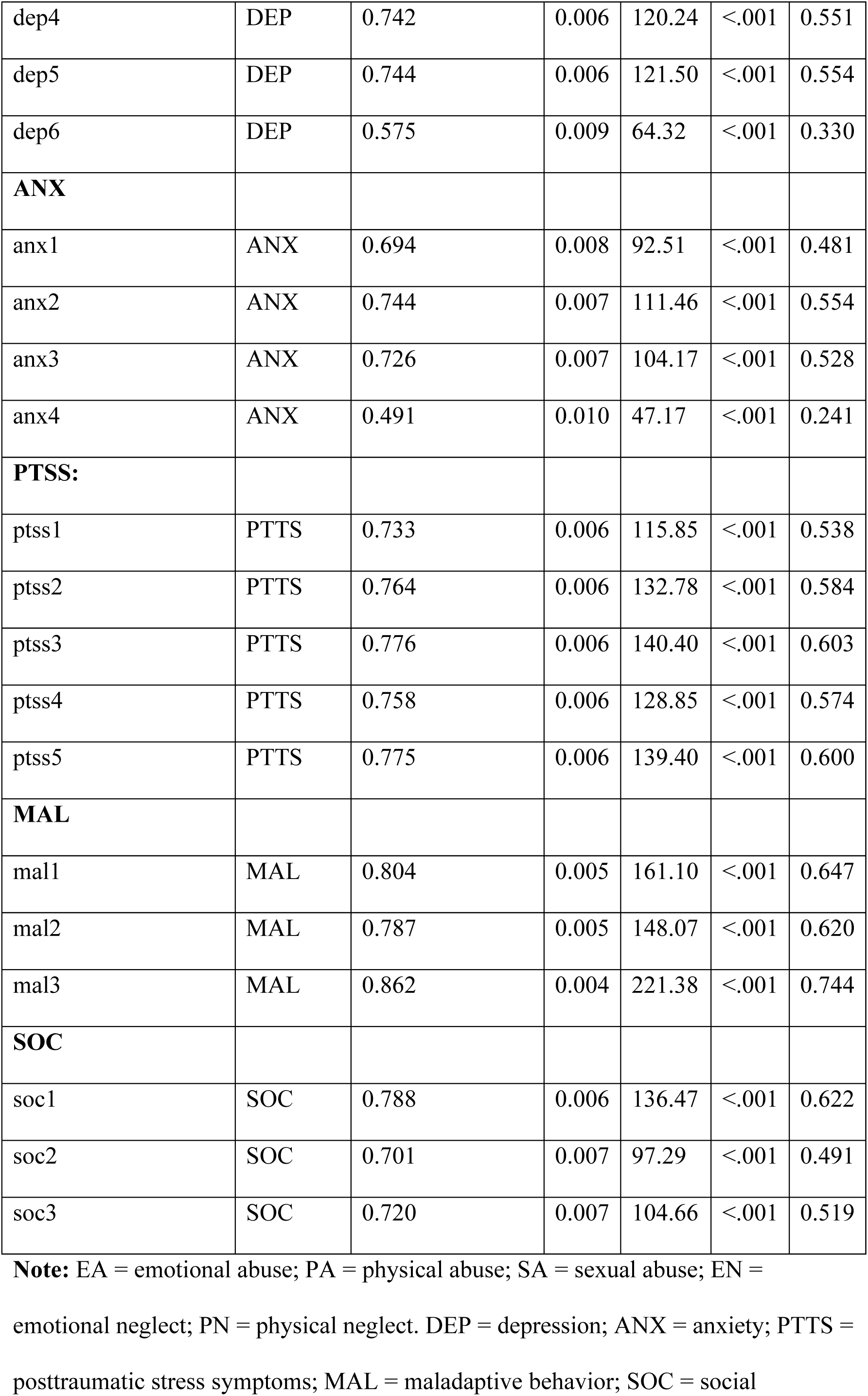
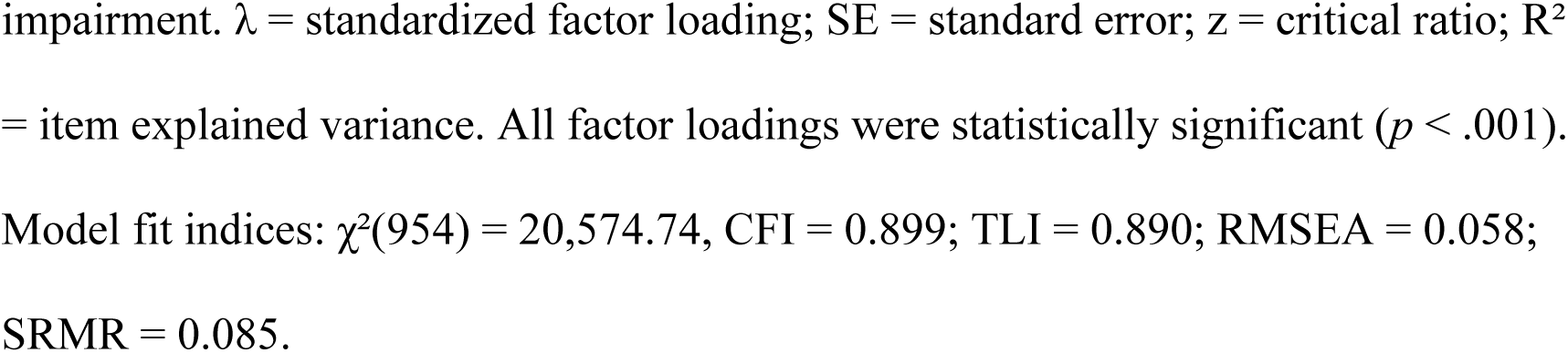
Confirmatory Factor Analysis (CFA) of the Childhood Trauma and Mental Health Outcome Questionnaire (N = 6,181)

As shown in Table 2, the measurement model demonstrated (χ²(954) = 20,574.74, CFI = 0.899, TLI = 0.890, RMSEA = 0.058, SRMR = 0.085).

Four items (soc4, sa15, pn23, pn24) were removed due to poor psychometric performance. The standardized factor loadings for the retained indicators were all statistically significant (*p* < .001) and ranged from 0.491 to 0.932, indicating strong relationships between observed indicators and their respective latent constructs. The removed items showed weaker factor performance, and their exclusion improved model adequacy. However, these modifications were interpreted cautiously because item performance may have been influenced by cultural interpretation or reverse-wording effects rather than solely by the underlying construct.

### 3.3 Measurement Invariance

Measurement invariance across gender and country groups was assessed using MGCFA, with results summarized in Table 3. The analysis supported configural, metric, and scalar invariance across gender, with minimal changes in model fit (ΔCFI ≤ 0.005). This indicates that the measurement structure functioned equivalently for male and female participants, allowing meaningful comparisons of latent constructs between genders.

**Table 3.** Measurement Invariance across Genders and Countries (N = 6,181)

| **Invariance Type/Model** | $\chi^2$ | df | CFI | ΔCFI | RMSEA | ΔRMSEA | Decision |
| --- | --- | --- | --- | --- | --- | --- | --- |
| **A. Gender Invariance** |  |  |  |  |  |  |  |
| Configural Model | 16,594.05 | 1,696 | 0.922 | — | 0.053 | — | Supported |
| Metric Model | 16,738.93 | 1,735 | 0.920 | 0.002 | 0.053 | 0.000 | Supported |
| Scalar Model | 17,094.03 | 1,774 | 0.915 | 0.005 | 0.054 | 0.001 | Supported |
| **B. Country Invariance** |  |  |  |  |  |  |  |
| Configural Model | 20,851.99 | 1,832 | 0.901 | — | 0.058 | — | Supported |
| Strict Metric Invariance | 21,260.10 | 1,869 | 0.899 | 0.002 | 0.058 | 0.000 | Rejected (statistically) |
| Partial Metric Invariance | 21,247.10 | 1,867 | 0.899 | 0.002 | 0.058 | 0.000 | Supported (practically) |
| Partial Scalar Invariance | 21,942.73 | 1,896 | 0.896 | 0.004 | 0.058 | 0.001 | Rejected (statistically; identification) |
| (Refined) |  |  |  |  |  |  | n issues) |

**Note:** CFI = Comparative Fit Index; RMSEA = Root Mean Square Error of
Approximation; $\Delta$ indicates change relative to the preceding model.
Gender invariance: fully configural, metric, and scalar invariance supported.
Country invariance: configural and partial metric invariance achieved; scalar
invariance not fully supported, indicating some cross-country differences in
intercepts.

Across the 17 participating countries, the configural invariance model demonstrated acceptable fit (CFI = 0.901; RMSEA = 0.058). Although strict metric invariance was statistically rejected, a partial metric invariance model was supported (ΔCFI = 0.002). The partial metric invariance model indicates that most factor loadings were comparable across countries, while a limited number of indicators demonstrated differential loading patterns. These differences suggest that some trauma indicators may be interpreted differently across cultural contexts; however, the majority of factor loadings remained invariant, supporting the validity of cross-national comparisons of structural associations.

### 3.4 Descriptive Statistics and Correlations

Following item refinement, the scales demonstrated good to excellent internal consistency, with Cronbach’s α values ranging from 0.755 (Anxiety) to 0.933 (Emotional Neglect).

Descriptive statistics for childhood trauma domains and mental health outcomes are presented in Table 4. Among trauma domains, physical neglect (M = 3.04, SD = 1.16) and emotional neglect (M = 2.44, SD = 1.23) showed the highest mean scores. In contrast, sexual abuse reported the lowest level (M = 0.61, SD = 0.91). Pearson correlation coefficients among study variables are presented in Table 5. All active abuse domains demonstrated significant positive correlations with mental health outcomes. Emotional abuse exhibited the strongest associations, particularly with depressive symptoms (r = .62, p < .001) and PTSS (r = .58, p < .001). When neglect domains were scored such that higher values represented greater neglect exposure, emotional neglect and physical neglect demonstrated unexpected negative correlations with several psychological outcomes. For example, emotional neglect was negatively correlated with depression (r = −.17, p < .001), PTSS (r = −.18, p < .001), and social impairment (r = −.32, p < .001). These associations were therefore not interpreted as simple reverse-scoring artifacts and were considered potential indicators of differential item functioning, cultural interpretation, or measurement characteristics requiring cautious interpretation.

**Table 4.** Descriptive Statistics, Internal Consistency Reliability, and Normality Indices for Childhood Trauma and Mental Health Outcome Questionnaire (N = 6,181)

| Subscale | Items | Cronbach's<br>$\alpha$ | McDonald's<br>$\omega$ | Mean<br>(SD) | Skewness | Kurtosis |
| --- | --- | --- | --- | --- | --- | --- |
| **Childhood**<br>**Trauma** |  |  |  |  |  |  |
| EA | 5 | 0.874 | 0.865 | 0.95<br>(0.97) | 1.12 | 0.54 |
| PA | 5 | 0.891 | 0.887 | 0.66<br>(0.93) | 1.67 | 2.12 |
| SA | 4† | 0.873 | 0.875 | 0.61<br>(0.91) | 1.78 | 2.65 |
| EN | 5 | 0.933 | 0.931 | 2.44<br>(1.23) | -0.42 | -0.95 |
| PN | 3† | 0.886 | 0.893 | 3.04<br>(1.16) | -1.26 | 0.62 |
| **Mental**<br>**Health**<br>**Outcomes** |  |  |  |  |  |  |
| DEP | 6 | 0.883 | 0.923 | 1.45<br>(0.97) | 0.49 | -0.44 |
| ANX | 4 | 0.755 | 0.877 | 1.33<br>(0.94) | 0.58 | -0.26 |
| PTTS | 5 | 0.874 | 0.916 | 1.39 | 0.57 | -0.53 |
|  |  |  |  | (1.05) |  |  |
| MAL | 3 | 0.859 | 0.893 | 1.39<br>(1.13) | 0.59 | -0.57 |
| SOC | 3† | 0.783 | 0.894 | 1.42<br>(1.11) | 0.54 | -0.62 |

**Notes:** EA = emotional abuse; PA = physical abuse; SA = sexual abuse; EN = emotional neglect; PN = physical neglect; DEP = depression; ANX = anxiety; PTTS = posttraumatic stress symptoms; MAL = maladaptive behavior; SOC = social impairment. † Items sa15, pn23, pn24, and soc4 were excluded prior to computation due to suboptimal psychometric performance. Cronbach's $\alpha$ values reflect internal consistency after item removal. Higher scores indicate greater severity or frequency of trauma exposure or psychological symptoms.

**Table 5.**
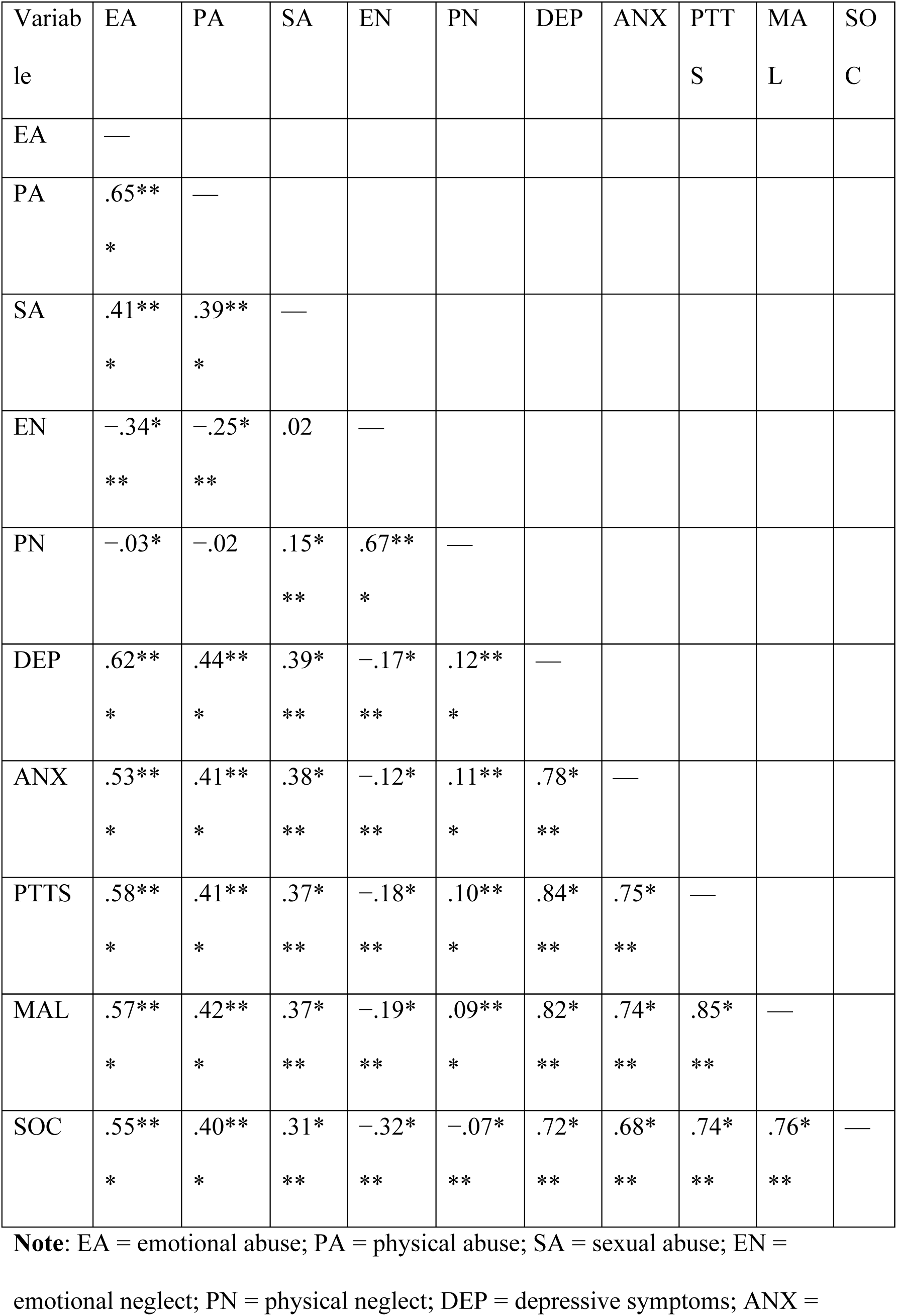
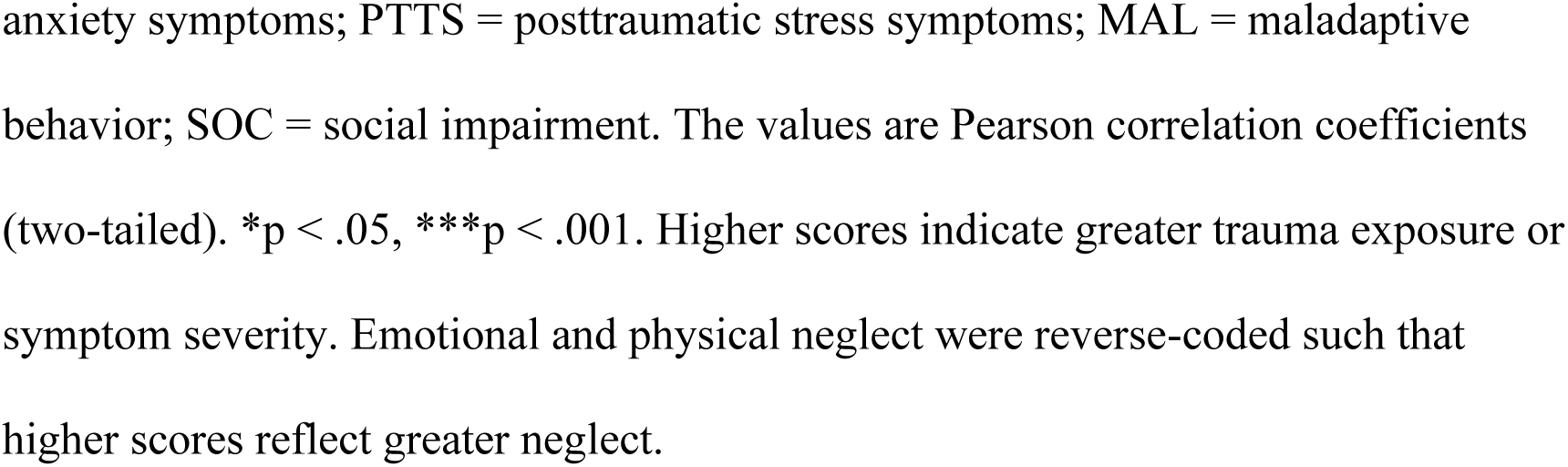
Pearson correlations between childhood trauma and mental health outcomes (N = 6,181)

### 3.5 Structural Equation Modeling and Regression Analysis

Multiple regression analyses controlling for age, gender, and education were conducted to identify predictors of mental health outcomes (Table 6). Emotional abuse emerged as the strongest independent predictor of depressive symptoms (β = 0.472, p < .001) and PTSS (β = 0.439, p < .001).

**Table 6.**
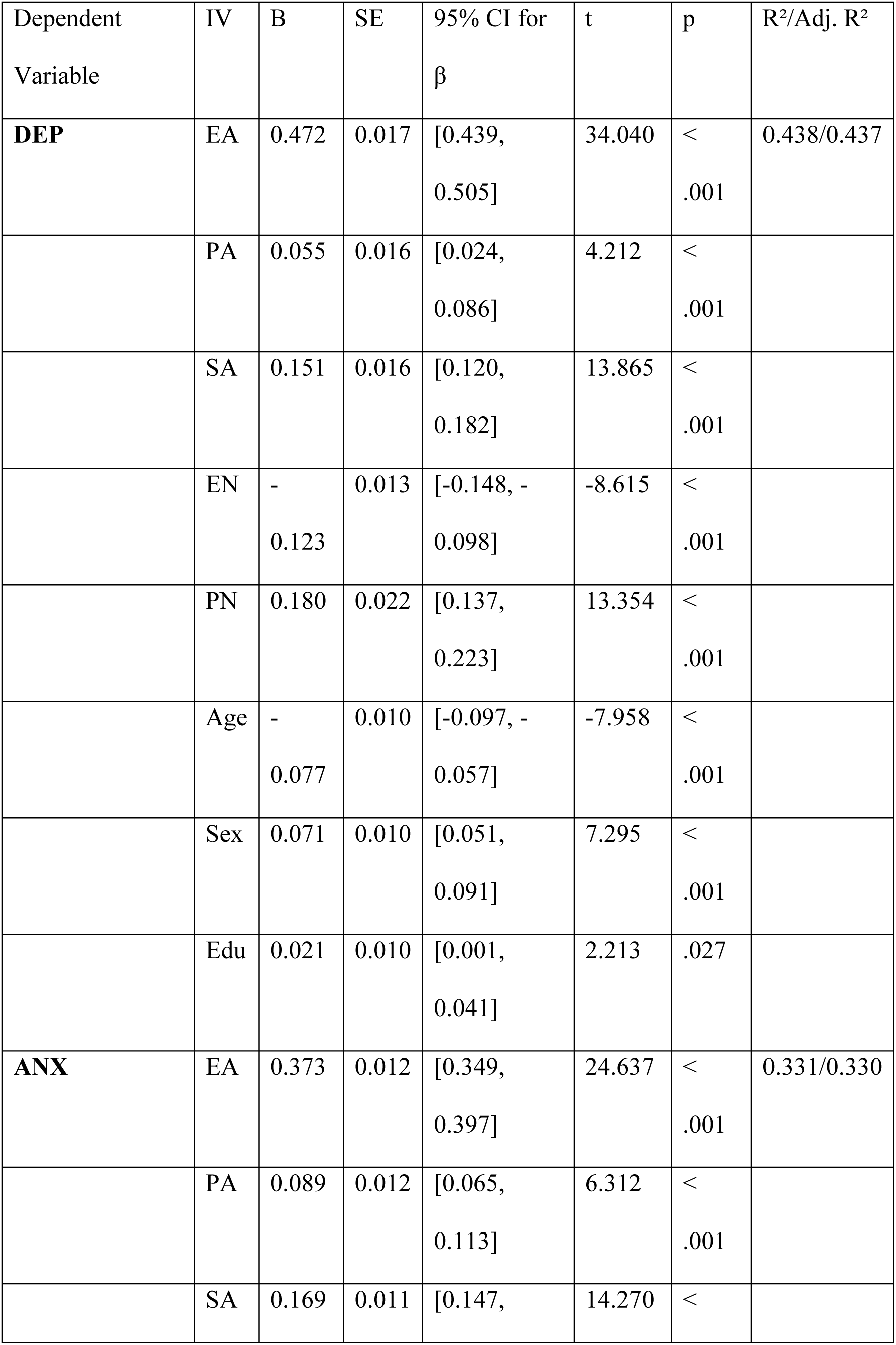
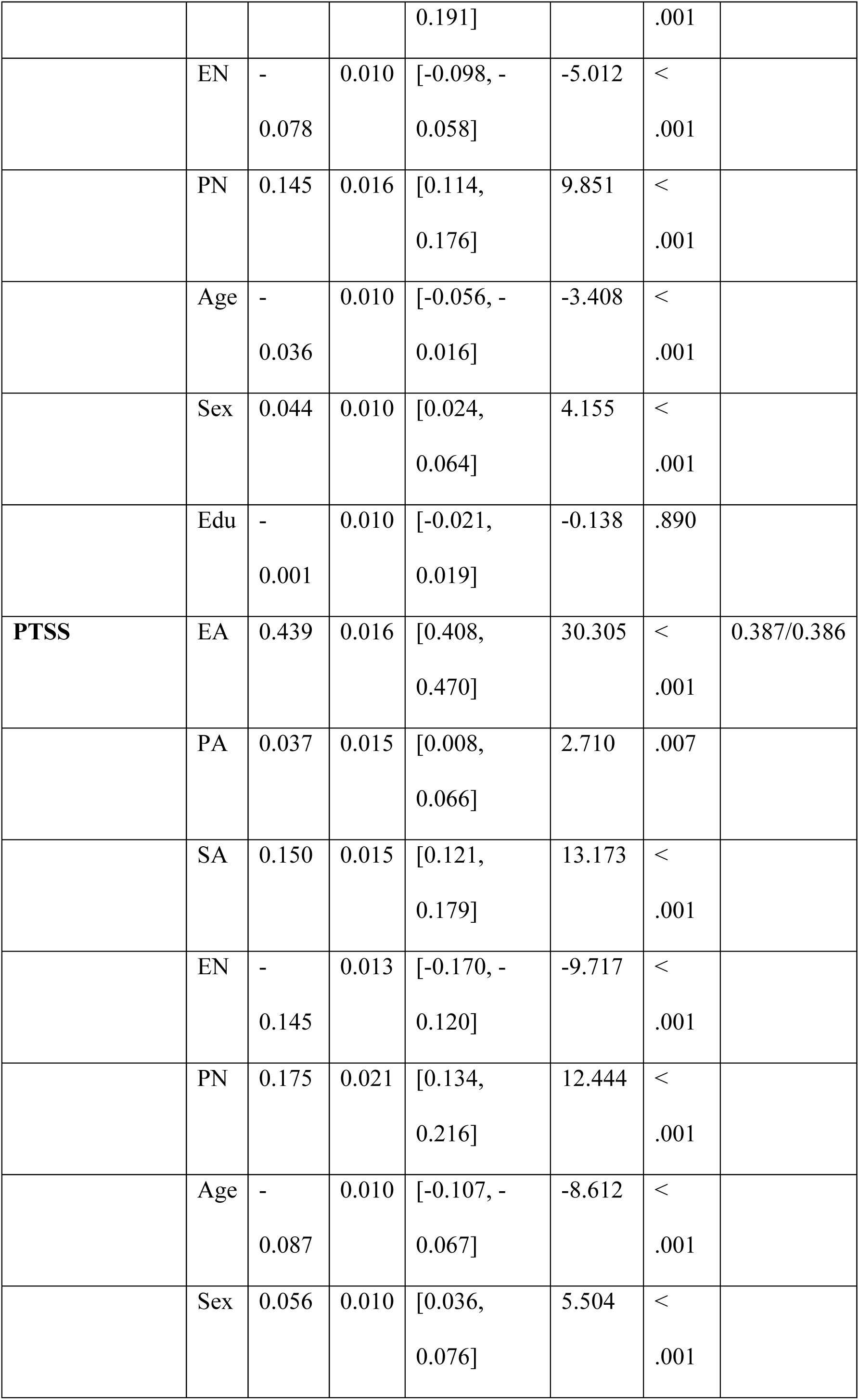
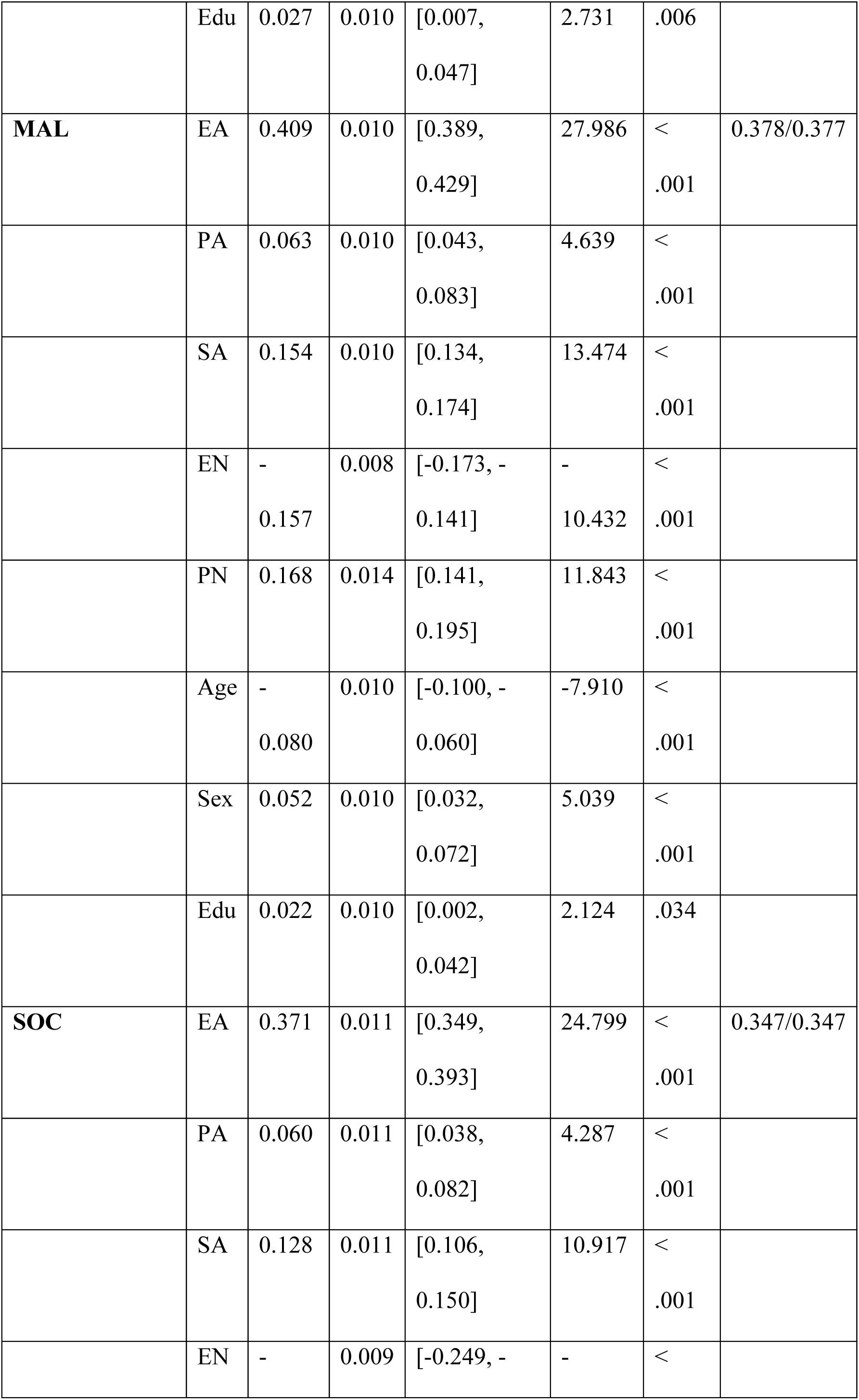
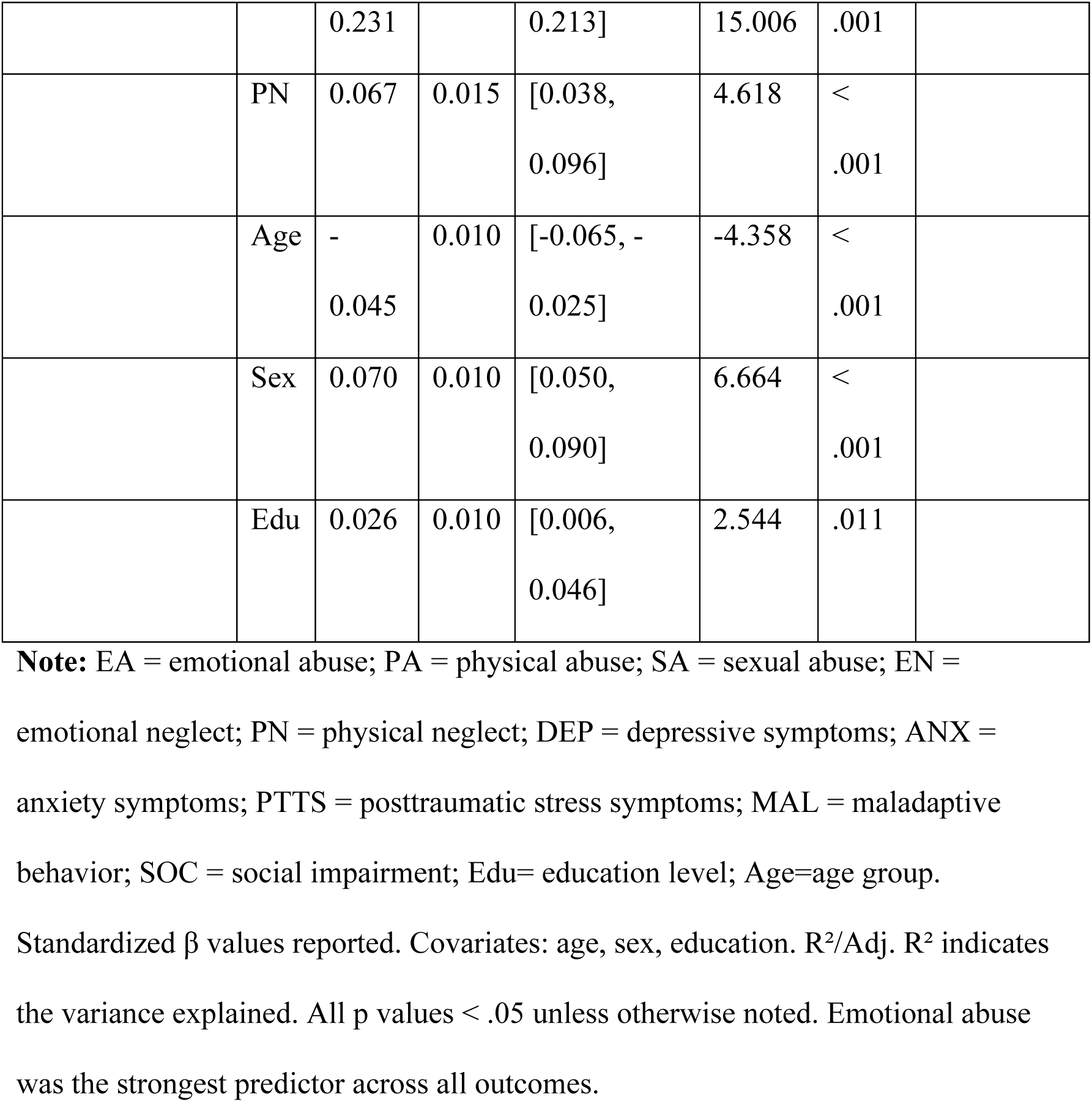
Multiple Regression Models Predicting Mental Health Outcomes from Childhood Trauma and Covariates (N=6181).

To account for trauma comorbidity, a second-order structural equation model (SEM) was estimated to examine whether a second-order latent construct predicted transdiagnostic mental health outcomes (Table 7; Figure 1). Within this higher-order model, emotional abuse was the strongest indicator of the global trauma factor (β = .942), followed by physical abuse (β = .758) and sexual abuse (β = .573).

**Figure 1.**
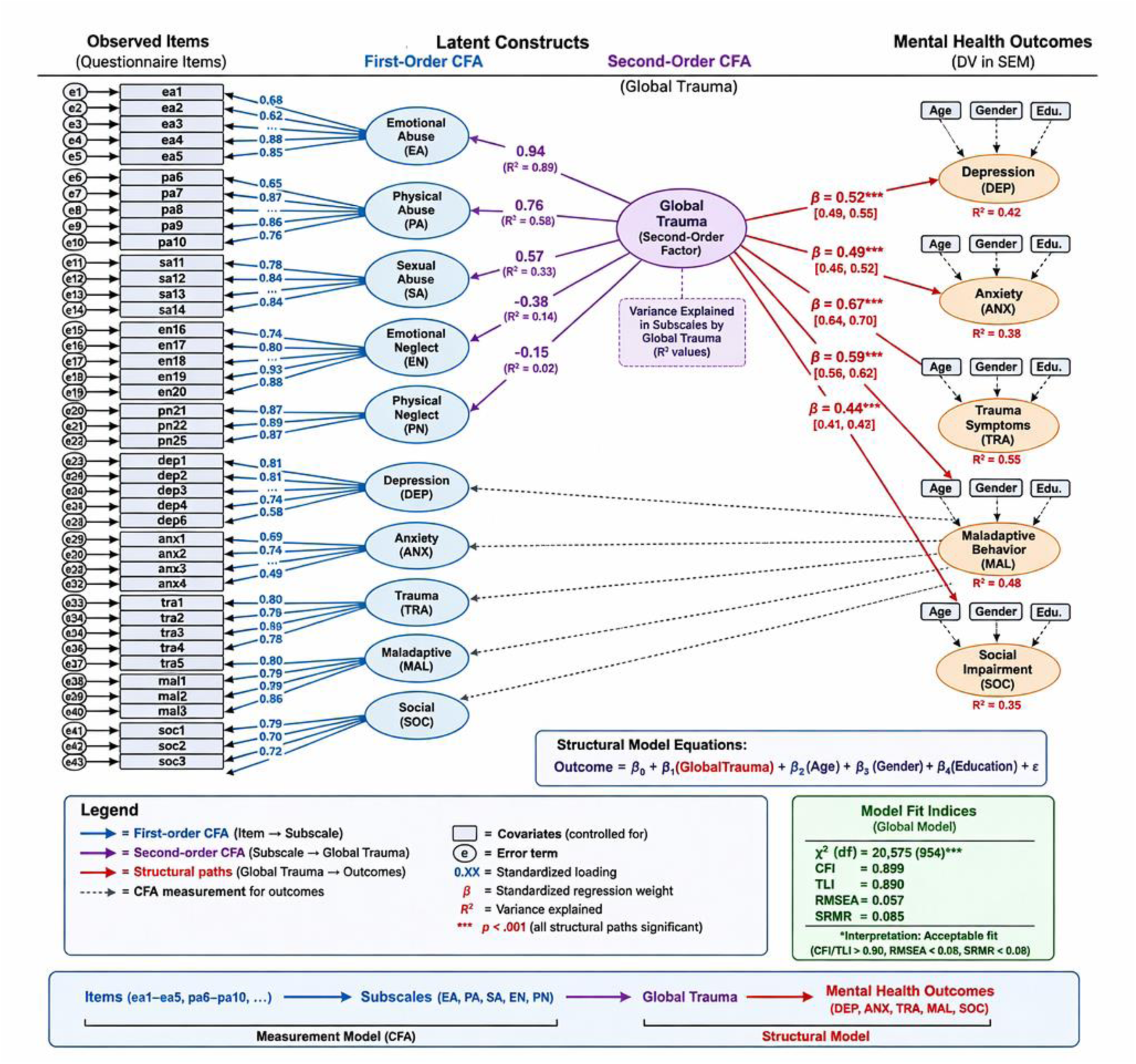
Second-order structural equation model (SEM) linking the latent global trauma construct to mental health outcomes (N = 6,181). *Note.* EA = emotional abuse; PA = physical abuse; SA = sexual abuse; EN = emotional neglect; PN = physical neglect; DEP = depressive symptoms; ANX = anxiety symptoms; TRA = trauma-related symptoms or PTTS = posttraumatic stress symptoms; MAL = maladaptive behavior; SOC = social impairment.

**Table 7.** Structural Equation Model: Global Childhood Trauma Predicting Mental Health Outcomes (N = 6,181).

| Path/Construct | $\beta$ | SE | 95% CI | z | p | R<sup>2</sup> |
| --- | --- | --- | --- | --- | --- | --- |
| **Second-Order Factor Loadings**<br>**(Global Trauma)** |  |  |  |  |  |  |
| EA | .942 | — | — | — | < .001 | .888 |
| PA | .758 | 0.016 | [.727, .789] | 37.53 | < .001 | .575 |
| SA | .573 | 0.020 | [.540, .606] | 34.04 | < .001 | .328 |
| EN | −.377 | 0.021 | [−.407, −.347] | −24.76 | < .001 | .142 |
| PN | −.152 | 0.021 | [−.181, −.123] | −10.43 | < .001 | .023 |
| **Structural Paths (Mental Health Outcomes)** |  |  |  |  |  |  |
| Global Trauma → DEP | .702 | 0.022 | [.668, .736] | 40.41 | < .001 | .515 |
| Global Trauma → ANX | .644 | 0.021 | [.607, .681] | 34.35 | < .001 | .427 |
| Global Trauma → PTSS | .679 | 0.022 | [.644, .714] | 37.58 | < .001 | .479 |
| Global Trauma → MAL | .681 | 0.024 | [.647, .715] | 39.06 | < .001 | .478 |
| Global Trauma → SOC | .667 | 0.024 | [.632,<br>.702] | 37.39 | <<br>.001 | .462 |

**Note:** EA = emotional abuse; PA = physical abuse; SA = sexual abuse; EN = emotional neglect; PN = physical neglect; DEP = depressive symptoms; ANX = anxiety symptoms; PTTS = posttraumatic stress symptoms; MAL = maladaptive behavior; SOC = social impairment. $\beta$ = standardized coefficient; SE = standard error; $R^2$ = variance explained. Emotional abuse loading was fixed to 1.0 to scale the latent construct. Global trauma significantly predicted all mental health outcomes ( $p <$ .001). Model fit indices: $\chi^2(954) = 20,574.74$ , CFI = 0.899; TLI = 0.890; RMSEA = 0.058; SRMR = 0.085.

The very high loading of emotional abuse suggests that this domain contributed substantially to the definition of the global trauma construct. Nevertheless, the remaining trauma domains retained significant contributions, supporting the multidimensional conceptualization of childhood adversity. A bifactor specification could further separate general trauma burden from domain-specific effects; however, the current analysis focused on evaluating the theoretically proposed second-order trauma framework.

Notably, emotional neglect and physical neglect showed negative loadings on the global trauma factor (β = −.377 and β = −.152, respectively). This unexpected pattern was identified as an important measurement finding rather than interpreted as evidence of a protective effect of neglect. The negative loadings may reflect differences in item functioning, cultural interpretation, or response patterns related to neglect domains.

The second-order latent factor significantly predicted all mental health outcomes (p < .001). The strongest effect was observed for depressive symptoms (β = .702), explaining 51.5% of the variance (R² = .515). Significant effects were also found for PTSS (β = .679; R² = .479), maladaptive behavior (β = .681; R² = .478), social impairment (β = .667; R² = .462), and anxiety (β = .644; R² = .427).

### 3.6 Group Comparisons

Results of demographic group comparisons are presented in Table 8. Independent sample t-tests indicated that females reported significantly higher levels of depressive symptoms than males (*t* = −10.586, *p* < .001, Cohen’s *d* = 0.29), as illustrated in Figure 2.

**Figure 2.**
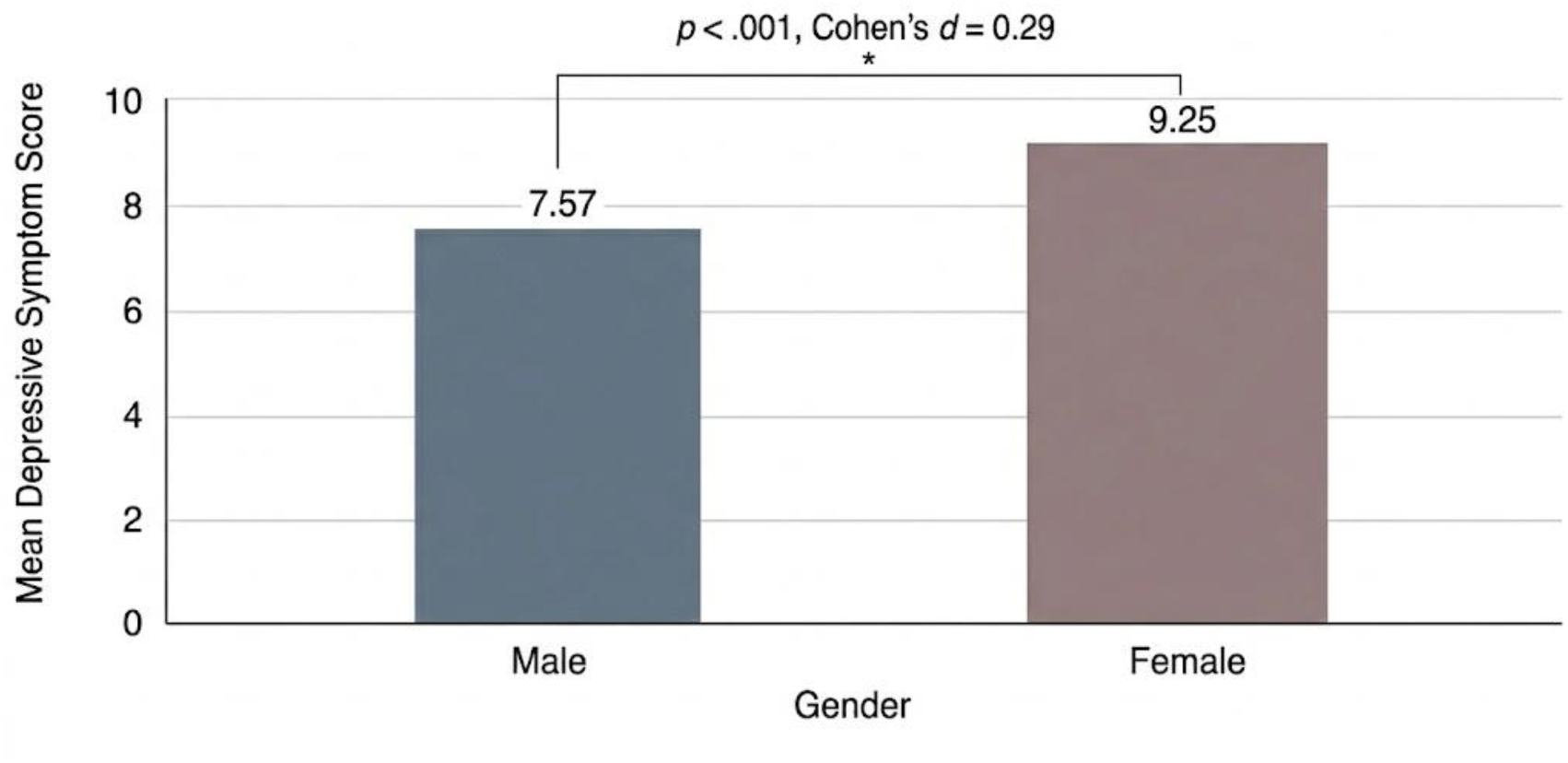
Mean depressive symptom scores by sex among study participants (N = 6,181). Note: Bar plots show the mean depressive symptom scores (6-item scale). Females reported significantly more symptoms than males did (t = -10.586, p < .001, Cohen’s d = 0.29). The error bars represent the standard error of the mean.

**Table 8.** Group Comparisons of Mental Health Outcomes (t test/ANOVA; N=6181).

| OV | GV | F/t | df | p | $\eta^2$ | Post hoc Summary |
| --- | --- | --- | --- | --- | --- | --- |
| DEP | Sex | t = -<br>10.586 | 6179 | <<br>.001 | 0.018 | Females (M = 9.25) > Males (M = 7.57); Cohen's d = 0.29.<br>Females reported significantly higher depressive symptoms. |
| ANX | Age | F =<br>14.258 | 3,<br>6177 | <<br>.001 | 0.007 | 18–25 > 26–35 > 36–45 > 46–55. Significant differences across age groups (Bonferroni/Tukey post hoc). |
| TRA | Edu | F =<br>3.944 | 4,<br>6175 | .003 | 0.003 | Bachelor > Diploma ≈ High school ≈ Postgraduate ≈ Uneducated. Some pairwise differences not significant; higher symptoms among lower education levels. |
| MAL | Occupation | F =<br>15.663 | 6,<br>6174 | <<br>.001 | 0.015 | Students & Health practitioners > Retired, Public sector, Freelancer, Private sector, Unemployed. Unemployed participants showed higher impairment. |
| SOC | Country | F =<br>14.332 | 16,<br>6164 | <<br>.001 | 0.036 | Highest scores: Iraq, Algeria, Jordan, Libya, Egypt; Lowest scores: UAE, Bahrain, Oman.<br>(Bonferroni/Tukey<br>comparisons). |

**Note:** OV = outcome variable; GV = grouping variable; DEP = depression; ANX = anxiety; PTTS = posttraumatic stress symptoms; MAL = maladaptive behavior; SOC = social impairment; Edu= education level; Age=age group. Effect sizes reported where appropriate (Cohen's d or $\eta^2$ ). Post hoc comparisons were performed via Bonferroni/Tukey tests.

Analysis of variance (ANOVA) revealed that anxiety symptoms varied significantly across age groups (*F* = 14.258, *p* < .001), with the highest levels observed among participants aged 18–25 years.

Additionally, social and functional impairment differed significantly across countries (*F* = 14.332, *p* < .001). Higher levels of impairment were reported among participants from Iraq, Algeria, Jordan, Libya, and Egypt, whereas lower levels were observed among respondents from the United Arab Emirates, Bahrain, and Oman.

## 4. Discussion

This study represents one of the largest cross-national investigations of childhood trauma and transdiagnostic psychopathology in the Arab world. Results indicate that cumulative childhood trauma is a strong predictor of adult mental health symptoms. The study findings demonstrate substantial explanatory associations, accounting for a considerable proportion of variance in adult depression (51.5%), PTSS (47.9%), and maladaptive behaviors (47.8%); however, these findings should be interpreted as associations rather than evidence of causal pathways due to the cross-sectional design. These effect sizes align with global meta-analyses identifying ACEs as a primary, preventable driver of mental illness across the lifespan.^31^

A key contribution of this study is validating the global trauma burden construct. The structural model confirms that emotional, physical, and sexual maltreatment are not merely co-occurring events but also cluster into a unified burden of adversity. This finding validates contemporary transdiagnostic models, suggesting that diverse forms of early trauma are damaged through shared neurobiological and psychosocial pathways rather than through isolated mechanisms.^10^

### 4.1 The Primacy of Emotional Abuse

A central finding of this investigation is the dominance of emotional abuse as the core component of the trauma experience. Emotional abuse exhibited the strongest loading on the global trauma burden (**λ**= .942) and consistently emerged in regression models as the most powerful independent predictor of depression, anxiety, and maladaptive behaviors. This suggests that psychological maltreatment—characterized by insults, humiliation, and terrorizing—constitutes the “core variance” of trauma in this population.

Unlike episodic physical abuse, emotional abuse often involves chronic interpersonal invalidation within the family. This sustained psychological erosion fundamentally undermines self-worth and disrupts emotional regulation development.^32–33^ Our findings are consistent with neurobiological research suggesting that emotional maltreatment may affect brain systems involved in salience processing and emotional regulation, including the amygdala and medial prefrontal cortex; however, because this study did not include neurobiological measurements, these mechanisms should be interpreted as plausible explanatory pathways rather than direct evidence from the current sample.^40^

### 4.2 Methodological Insights: The Paradox of Neglect

While emotional and physical neglect were the most frequently endorsed forms of adversity, they presented negative loadings on the global trauma burden and inverse associations with active abuse types in our multivariate models. This apparent paradox may partially reflect methodological artifacts associated with reverse-coded CTQ-SF items (e.g., “I felt loved,” “I had enough to eat”) rather than solely genuine psychological phenomena.

In multilingual and cross-cultural survey contexts, reverse-coded items may introduce systematic measurement error, including acquiescence response bias and comprehension difficulties during translation. Consequently, the unusual negative loadings observed for emotional and physical neglect may reflect a wording-related “method factor,” whereby covariance is influenced by item phrasing rather than the underlying trauma construct itself. Nevertheless, sociocultural factors may also contribute to these findings. In the Arab context, where family cohesion is strongly emphasized, respondents may simultaneously report high levels of familial “love” and disciplinary “abuse.” Furthermore, the Threat versus Deprivation model suggests that abuse and neglect influence neurodevelopment differently: abuse sensitizes threat detection systems associated with anxiety and PTSD, whereas deprivation may disrupt reward processing and contribute to emotional blunting rather than overt distress symptoms. ^34, 35, 41^

Taken together, these findings highlight the importance of cautious interpretation of neglect subscales within cross-cultural research settings. Future studies may benefit from bifactor approaches or explicit method-factor modeling to isolate variance associated with reverse-worded items and further refine the latent structure of global trauma.

### 4.3 Sociocultural Context and Measurement Validity

The interpretation of these results must be grounded in the sociocultural fabric of the MENA region. The rigorous establishment of measurement invariance across genders and countries is a critical methodological advance of this study. This finding confirms that the constructs of trauma and mental health are measured equivalently across diverse Arab nations—from the Gulf to the Maghreb—validating the use of these tools for cross-national comparison.

However, the low reported prevalence of sexual abuse likely reflects deep-seated cultural taboos and fear of stigma, suggesting underreporting.^36, 37^ Additionally, group comparisons revealed that participants from conflict-affected nations (Iraq, Libya, Syria, and Yemen) reported the highest levels of social and functional impairment. This finding supports the “double burden” hypothesis, where the intrafamilial trauma of childhood is compounded by the macrolevel stressors of political instability, displacement, and structural violence, severely impeding functional recovery.^38^

### 4.4 Neurobiological implications

The strong associations found between the global trauma burden and transdiagnostic symptoms support the “toxic stress” model. Chronic activation of the stress response system during sensitive developmental windows is known to dysregulate the hypothalamic–pituitary–adrenal (HPA) axis.^39^ Although the present study did not directly measure neuroendocrine or neural markers, the broad pattern of associations across mood, anxiety, and behavioral outcomes is compatible with previous neurobiological models proposing that early adversity may contribute to generalized vulnerability through stress-response dysregulation and related developmental mechanisms.

### 4.5 Strengths and Limitations

The strengths of this study include its large multinational sample, the use of advanced structural modeling (SEM/CFA), and the rigorous testing of measurement invariance. However, limitations must be noted. First, the cross-sectional design precludes causal inference. Second, the reliance on self-report measures introduces recall bias. Specifically, individuals currently experiencing severe psychological distress may recall past events more negatively—a phenomenon known as mood-congruent memory bias—which could artificially inflate the magnitude of the observed associations (e.g., the strong predictive paths to depression and trauma-related symptoms). Third, the sample was recruited via social media, resulting in an overrepresentation of younger, educated females; consequently, the findings may not be fully generalizable to older, rural, or offline populations. Fourth, the study did not account for several critical unmeasured confounders. Variables such as socioeconomic status (SES), current acute life stressors (including ongoing war exposure and financial instability), family psychiatric history, and access to mental health services are likely associated with both the likelihood of trauma exposure and the severity of adult psychopathology. Finally, specific items (e.g., soc4, sa15) were excluded to achieve model fit, indicating a need for further psychometric adaptation of these instruments for Arabic-speaking populations.

Furthermore, although the magnitude of variance explained in this study was substantial, direct comparison with Western studies is challenging because previous investigations have used heterogeneous trauma measures, outcomes, samples, and analytical approaches. Nevertheless, the observed associations are broadly consistent with Western literature demonstrating robust relationships between childhood adversity and later psychopathology, while the present findings extend this evidence to a large Arab multinational sample.

### 4.6 Conclusion

Childhood trauma is a pervasive public health crisis worldwide. The “Global Trauma” burden—driven primarily by emotional abuse—predicts a wide array of adult psychiatric morbidities. Given the observational cross-sectional design, these findings do not establish that trauma directly causes later psychopathology; rather, they highlight the importance of considering childhood adversity history when assessing and supporting individuals with mental health difficulties. Public health policies should prioritize trauma-informed care and parenting programs targeting emotional maltreatment.

## Author’s Contribution Statement

*Conceptualization*: K.M.A., B.A.O.; *Methodology*: K.M.A., B.A.O.; *Data Curation*: D.M.E., Y.E.H.; *Software & Validation*: A.N.A., S.E.F.; *Investigation & Analysis*: M.S.A., B.H.H., A.S.A., K.M.A.; *Visualization*: A.S.A., Y.N.R.; *Writing – Original Draft*: K.M.A.; *Writing – Review & Editing*: M.W.S., L.S.K., A.A.A., N.A.Q.; *Resources*: A.A.A., N.A.Q.; *Supervision*: K.M.A.; *Project Administration*: K.M.A.; *Funding Acquisition*: N/A.

## Ethical Considerations

This study was conducted in accordance with the ethical principles of the Declaration of Helsinki. Participation was voluntary, and electronic informed consent was obtained from all participants before accessing the questionnaire. No personally identifiable information was collected, and all responses were recorded anonymously. **Data availability**

The datasets generated and analyzed during the current study are available from the corresponding author upon reasonable request.

## Use of artificial intelligence (AI)

No generative artificial intelligence tools were used in data generation, statistical analysis, or interpretation.

## Acknowledgments

The authors would like to express their gratitude to **ReseMeds Academy** for providing the technical framework, statistical consulting, and research coordination that facilitated this multicountry study. Furthermore, the authors would like to thank all individuals who contributed to the data collection and coordination of this multicountry study. Their efforts in participant recruitment, survey administration, and field coordination were essential to the successful completion of this research. *Data Collectors:* Abdalrhman Ibrahim Ali Eisa; Abdullah Essam Hassan; Abdurrhman Elhadi Abdullah; Abeer Atia Edriwi; Ahmed Yacine Hadef; Alhanof Mabrook; Amal Fouad Mzayen; Amna Yaser Banihammad; Areena Abed; Asia Ahmed; Arowa Osman Atiya; Ayham R. Zidat; Dina Eldawoody; Doaa Arrahim; Effa Mohamed Osman Mohamed Abubaker; Eshrak Mohammed Alaboud; Esraa Magzoub Elrayah Mohamed Ahmed; Guitouni Yasmine; Haider Ali Raheem; Hamza Adel Mahmoud; Heba Mobark Ahmed; Hiba Hassan Matroud; Hind El Azzazi; Iman Al-Haddawi; Iman Mohammed Sulaiman Al Dhawyani; Jamil Jamal Mahmoud; Jokha Qasim Al Busaidi; Juhaina Nafea Mohammed; Kareem Muneer Hasan; Karrar Mushtaq Talib; Khalid Rajab Mukhtar; Lara Khaled Barbar; Maad Nasser Mohamed; Madleen Algadsi; Mahmoud Mohamed Al-Ashram; Manar Moamer Sahloul; Maryam A. Ameer Emam; Meddah Sara; Meiad Awad Elsheikhidris Abdelrahman; Moamen Haitham Ramadan; Mostafa Houssam Alwan; Mustafa Raed Naeem; Munerah N A Alrashidi; Najwa Abdalhady Nasr; Najwa Mahmoud Kouli; Nazar Yousif Yahya; Noor Badr Al-Raai; Rahaf Khaldoun Al-Qawasmi; Rahaf Mohamad Hamza; Reem Ibrahim Mohamed; Reem Salman Almahari; Roua Houssam Alwan; Sana Adel Badra; Sara Taieb Jafleh; Sawsan Jaffar Almadhoob; Wafa Bakhit Ibrahim Salmah; Walaa Althabati; Wasim Akram Abu Khousa; Zaina Husain Traif; Zainab Nasser Hassan.

## Appendix A

**Survey Questionnaire (Excerpt)**

### Section One: Demographic Information

- **Gender:** Male / Female
- **Age:** 18–25 / 26–35 / 36–45 / 46–55 years
- **Education:** Uneducated / High school / Diploma degree / Bachelor’s degree / Postgraduate degree
- **Occupation:** Student / Unemployed / Freelancer / Retired / Private sector / Public sector / Health practitioner
- **Country:** Saudi Arabia, UAE, Bahrain, Oman, Qatar, Jordan, Iraq, Palestine, Lebanon, Syria, Egypt, Sudan, Yemen, Morocco, Algeria, Tunisia, Libya

### Section Two: Childhood Trauma

Please indicate the extent to which you agree with the following statements, using the scale below:

(0) Never true (1) Rarely true (2) Sometimes true (3) Often true (4) Very often true

### Emotional Abuse Experiences

1. Did your family members call you names like “stupid,” “lazy,” or “ugly”?
2. Did you believe that your parents wished you had never been born?
3. Did you feel that a member of your family hated you?
4. Did your family members say hurtful or humiliating things to you?
5. Do you believe that you were subjected to emotional (psychological) abuse?

### Physical Abuse Experiences

6. Were you hit so hard by a family member that it required a visit to a doctor or hospital?
7. Did your family members hit you severely enough to leave bruises or marks?
8. Were you punished with hard objects like a belt or a cord?
9. Do you believe that you were subjected to physical abuse?
10. Did someone, like a teacher or a neighbor, notice signs of being hit on you?

### Sexual Abuse Experiences

11. Did anyone try to touch you in an inappropriate way or make you touch them?
12. Did you feel forced to do things of a personal nature against your will?
13. Were you pressured to participate in inappropriate activities?
14. Did you face disturbing behaviors of a private nature?
15. Did you feel safe from inappropriate behaviors during your childhood?

### Emotional Neglect Experiences

16. Was there someone in your family who made you feel important? (*Reverse scored*)
17. Did you feel loved in your family? (*Reverse scored*)
18. Did your family members care for one another? (*Reverse scored*)
19. Were your family members emotionally close? (*Reverse scored*)
20. Was your family a source of support for you? (*Reverse scored*)

### Physical Neglect Experiences

21. Did you have enough to eat? (*Reverse scored*)
22. Was there someone to take care of you and protect you? (*Reverse scored*)
23. Were your parents under the influence of alcohol or drugs to the extent that they neglected the family?
24. Did you have to wear dirty clothes?
25. Was there someone to take you to the doctor when needed? (*Reverse scored*)
26. Was there something you wanted to change about your family?
27. Was your childhood perfect? (*Reverse scored*)
28. Was your family the best family in the world? (*Reverse scored*)

### Section Three: The Impact of Childhood Trauma on Mental Health

Please select the most appropriate answer for each statement using the scale below:

(0) Never (1) Rarely (2) Sometimes (3) Often (4) Very Often

### Depressive Symptom

1. How often do you suffer from depressive symptoms, such as sadness or loss of interest?
2. How often do you feel guilt or shame?
3. How often do you find it difficult to concentrate or make decisions?
4. How often do you feel hopeless about the future?
5. How often do you feel worthless or have low self-esteem?
6. How often do your appetite or weight change due to your psychological state?

### Anxiety Symptom

7. How often do you feel anxious or excessively worried?
8. How often do you suffer from physical symptoms (e.g., headaches, stomach problems) due to anxiety?
9. How often do you experience difficulty sleeping?
10. Do you experience recurrent worry?

### Posttraumatic Stress Symptoms (PTSD)

11. How often do you experience painful memories or flashbacks?
12. How often do you feel disconnected from reality?
13. Do you avoid situations that remind you of past traumas?
14. How often do you experience recurring intrusive thoughts?
15. How often do you suffer from emotional numbness or detachment?

### Maladaptive Behavior / Impulse-Control

16. How often do you engage in self-harm behaviors or have suicidal thoughts?
17. How often do you feel irritable or excessively angry?
18. How often do you resort to addiction or negative coping behaviors?

### Social & Functional Impairment

19. To what extent do psychological symptoms affect your daily life?
20. How often do you feel lonely or isolated?
21. Do you find it difficult to trust others or form relationships?
22. Do you generally feel happy and secure? (*Reverse scored*)

